# White matter connectivity disruptions in the pre-clinical continuum of psychosis: A connectome study

**DOI:** 10.1101/365064

**Authors:** Lena K. L. Oestreich, Roshini Randeniya, Marta I. Garrido

**Affiliations:** Queensland Brain Institute, The University of Queensland, Brisbane 4072, Australia; Centre for Advanced Imaging, The University of Queensland, Brisbane 4072, Australia; Australian Centre of Excellence for Integrative Brain Function, The University of Queensland, Brisbane 4072, Australia; School of Mathematics and Physics, The University of Queensland, Brisbane 4072, Australia

**Keywords:** structural connectome, diffusion MRI, connectivity, psychotic-like experiences, continuum of psychosis, psychosis biomarker

## Abstract

**Background:** Widespread white matter disruptions in schizophrenia have been commonly reported, but it remains unanswered whether these abnormalities are associated with schizophrenia specifically or whether they range along a psychotic continuum into the healthy population. Investigating the extent of white matter connectivity disruptions specific to psychotic-like experiences in healthy individuals is insofar important as it is a necessary first step towards the development of prodromal psychosis biomarkers.

**Methods:** High resolution, multi-shell diffusion-weighted magnetic resonance images were acquired from 89 healthy individuals. Whole-brain white matter fiber tracking was performed to quantify the strength of white matter connections. Network-based statistics were applied to white matter connections in a regression model in order to test for a linear relationship between streamline count and psychotic-like experiences.

**Results:** A significant subnetwork was identified whereby streamline count declined with increasing quantity of psychotic-like experiences. This network of significant connectivity reductions affected all cortical lobes, subcortical structures and the cerebellum.

**Conclusion:** A widespread network of linearly declining connectivity strength with increasing number of psychotic-like experiences was identified in healthy individuals. This finding is in line with white matter connectivity reductions reported from early to chronic stages of schizophrenia. We suggest that these white matter changes might be a potential biomarker for the identification of individuals at high risk for transitioning to psychosis.

## 1. Introduction

Schizophrenia has long been conceptualized as a disorder of dysconnectivity (Friston, 1998) whereby poorly integrated brain regions are thought to lead to impaired functioning and the development of psychotic symptoms such as hallucinations and delusions. Neuronal connections of the central nervous system, which constitute the human connectome, are highly interdependent and crucial for healthy brain functioning (Narr & Leaver, 2015). Dysconnectivity at any level of these networks may impact on other networks and affect behaviour and cognition in such a way that disorders like schizophrenia can manifest (Narr & Leaver, 2015). In support of this hypothesis, a multitude of studies have reported abnormal functional, structural and effective connectivity in patients with schizophrenia (Pettersson-Yeo, Allen, Benetti, McGuire, & Mechelli, 2011).

Diffusion-weighted imaging (DWI) studies using region-of-interest (ROI) approaches have reported heterogeneous findings of localized white matter disruptions in schizophrenia (Parnanzone et al., 2017). In recent years, brain connectivity studies in schizophrenia have benefited from advances in neuroimaging connectomics (Fornito, Zalesky, Pantelis, & Bullmore, 2012), which enables the mapping of large-scale structural and functional brain networks across the brain without the requirement for a-priori hypotheses about potentially affected ROIs (Sporns, 2011). Structural connectomics studies suggest that white matter pathology in schizophrenia is more widespread than previously reported in studies using ROI approaches, affecting all cerebral lobes, the cerebellum and encompassing most of the brain’s white matter (Di Biase et al., 2017; van den Heuvel, Mandl, Stam, Kahn, & Hulshoff Pol, 2010; Zalesky et al., 2011).

A further explanation as to why previous DWI studies in schizophrenia reported vastly inconsistent findings might be the fact that studies with chronic schizophrenia patients are often biased by confounding variables like medication, hospitalization, deterioration of cognitive functioning and comorbidities like substance abuse (Frith, 1992). It is therefore unclear whether observed white matter changes are due to the illness itself or a result of these confounding variables. A study investigating 22q11 deletion syndrome, which is a genetic risk factor for schizophrenia, observed reduced network coherence, while other graph indices were preserved, which indicates that disruptions in certain white matter networks are associated with a genetic vulnerability for schizophrenia (Ottet et al., 2013). Similar to findings in patients with schizophrenia, (Collin, Kahn, de Reus, Cahn, & van den Heuvel, 2014) reported reduced rich club connectivity in unaffected siblings of patients with schizophrenia, suggesting a familial vulnerability for schizophrenia. Lastly, Di Biase et al. (2017) reported localized white matter reductions in patients with recent-onset schizophrenia compared to widespread white matter disruptions in chronic schizophrenia patients. Importantly, Di Biase et al. (2017) reported a significant linear decline of white matter connectome microstructure from early to chronic schizophrenia, indicating that white matter pathology deteriorates with illness duration.

So far, only one study investigated structural connectome changes in healthy individuals with psychotic-like experiences (i.e. sub-clinical psychotic symptoms in the general population; Drakesmith et al., 2015). This is insofar important, as prodromal psychotic-like experiences have been implicated in the development of florid psychosis (Kelleher & Cannon, 2011; Poulton et al., 2000) and may therefore help to identify individuals at high-risk for developing psychosis, who might benefit from prophylactic treatments. The study by Drakesmith et al. (2015) did not find any significant brain network differences in the streamline count between the high and low psychotic experiences groups. However, the topology measures global and local efficiency as well as global density were found to be significantly reduced in healthy individuals with psychotic-like experiences. These findings were interpreted to indicate that white matter changes in healthy individuals with psychotic-like experiences are subtle but detectable when examining network topology (Drakesmith et al., 2015).

The study by Drakesmith et al. (2015) used the psychotic-like symptoms semi-structured interview, which only assesses the positive symptoms hallucinations, delusions and bizarre symptoms (Horwood et al., 2008). However, individuals at risk for developing psychosis also possess various other sub-clinical symptoms, such as depression, negative symptoms and distress (Yung et al., 2007). In order to get a comprehensive picture of prodromal psychotic-like experiences in psychologically healthy individuals, it is therefore necessary to assess a variety of sub-clinical symptoms in addition to positive symptoms.

The present study set out to investigate structural connectome changes in a group of healthy individuals with psychotic-like experiences as measured by the prodromal questionnaire, which is a self-report measure, including questions about positive, negative and depressive symptoms as well as distress associated with these symptoms, and general functioning (Loewy, Bearden, Johnson, Raine, & Cannon, 2005). It was hypothesised that as the number of psychotic-like experiences increases, white matter connectivity would decrease in the same brain networks previously found to decline with illness duration in schizophrenia (Di Biase et al., 2017).

## 2. Materials and Methods

### 2.1 Participants

Eighty-nine healthy participants were recruited through the online recruitment system SONA and the weekly electronic newsletter UQ Update, which is distributed to staff and alumni at the University of Queensland, Australia. This study was approved by the University of Queensland Research Ethics Committee. All participants gave written informed consent and were monetarily reimbursed for their time. Participants ranged in age from 18 – 63 years (*M* = 24.69, *SD* = 10.13), 92.1% (*n* = 82) reported to be right-handed and 51% (*n* = 57.3) were female. All participants completed the prodromal questionnaire (PQ), which is a 92-item self-report screening measure for prodromal psychotic symptoms (Loewy et al., 2005) and includes items to assess depressive, positive and negative symptoms as well as distress. All items are rated on a scale from 0-5, whereby 0 = never, 1 = 1-2times, 3 = once per week, 4 = few times per week and 5 = daily. Each question is followed-up with a question asking whether the experience has been distressing (yes/no).

### 2.2 Data acquisition

T1-weighted magnetic resonance imaging (MRI) scans were acquired on a Siemens Trio 3T system (Erlangen, Germany) with the magnetisation-prepared two rapid acquisition gradient echo (MP2RAGE) sequence (Marques et al., 2010) with field of view (FoV) 240mm, 176 slices, 0.9mm isotropic resolution, TR=4000ms, TE=2.92ms, TI1=700ms, TI2=2220ms, first flip angle=6°, second flip angle=7°, and 5min acquisition time. Two diffusion-weighted (DW) image series were acquired using an echo-planar imaging (EPI) sequence. The first DW image series was acquired with the following parameters: FoV 220mm, phase partial Fourier (PPF) 6/8, parallel acceleration factor 2, 55 slices, 2mm isotropic resolution, 33 diffusion-sensitization directions at *b*=1000s/mm^2^ and one *b*=0 volume, TR=8600ms, TE=116ms, and 5min acquisition time. The second DW image series had a FoV of 220mm, PPF 6/8, parallel acceleration factor 2, 55 slices, 2mm isotropic resolution, 64 diffusion-sensitization directions at *b*=3000s/mm^2^ and two *b*=0 volumes, TR=8600ms, TE=116ms, and 10min acquisition time. Three additional *b*=0 images were acquired interspersed between the DW image series and the MP2RAGE sequence, with reversal of the acquisition direction along the phase-encoded axis for one of the three images and acquisition time of 30ms each.

### 2.3 Pre-processing and connectome generation

Figure 1 summarises the analysis workflow. DW images were corrected for eddy current distortions and head movements using the FSL TOPUP (Smith et al., 2004) and EDDY (Andersson & Sotiropoulos, 2016) commands and signal intensity inhomogeneities were removed (Zhang, Brady, & Smith, 2001). The remaining processing steps were conducted in the MRtrix3 toolbox (Tournier, Calamante, & Connelly, 2012). T1- and DW-weighted images were co-registered using boundary-based registration (Greve & Fischl, 2009). A five-tissue-type segmented image (cortical grey matter, white matter, sub-cortical grey matter, cerebrospinal fluid, pathological tissue) was generated from the structural images pre-processed with the recon-all command in FreeSurfer (Dale, Fischl, & Sereno, 1999). Response functions were estimated using the multi-shell, multi-tissue algorithm implemented in MRtrix3 (Jeurissen, Tournier, Dhollander, Connelly, & Sijbers, 2014). Multi-tissue constrained spherical deconvolution was applied to obtain fiber orientation distributions (Jeurissen et al., 2014). Eighty-four structural connectome nodes were defined using the Desikan-Killiany cortical segmentation atlas (Desikan et al., 2006). Individual tractograms were generated for each participants using anatomically constrained tractography (Smith, Tournier, Calamante, & Connelly, 2012) with the 2^nd^ order integration over Fibre Orientation Distribution (iFOD2) algorithm (Tournier et al., 2012). Tractograms were generated until 100 million streamlines were obtained with a length of 5-250mm, step size of 1mm and FOD amplitude threshold of 0.1. The spherical-deconvolution informed filtering of tractograms (SIFT) algorithm was applied to reduce the overall streamline count to 10 million streamlines, which are more biologically meaningful (Smith, Tournier, Calamante, & Connelly, 2013). To generate individual connectivity matrices, the streamlines from the tractograms were then mapped onto the nodes of each participants’ parcellation image which was generated from the anatomical (T1-weighted) images. Separate connectivity matrices were populated with the streamline count between the corresponding pairs of nodes, as well as fractional anisotropy (FA) estimates averaged over all voxels traversed by streamlines between node pairs.

**Figure 1.**
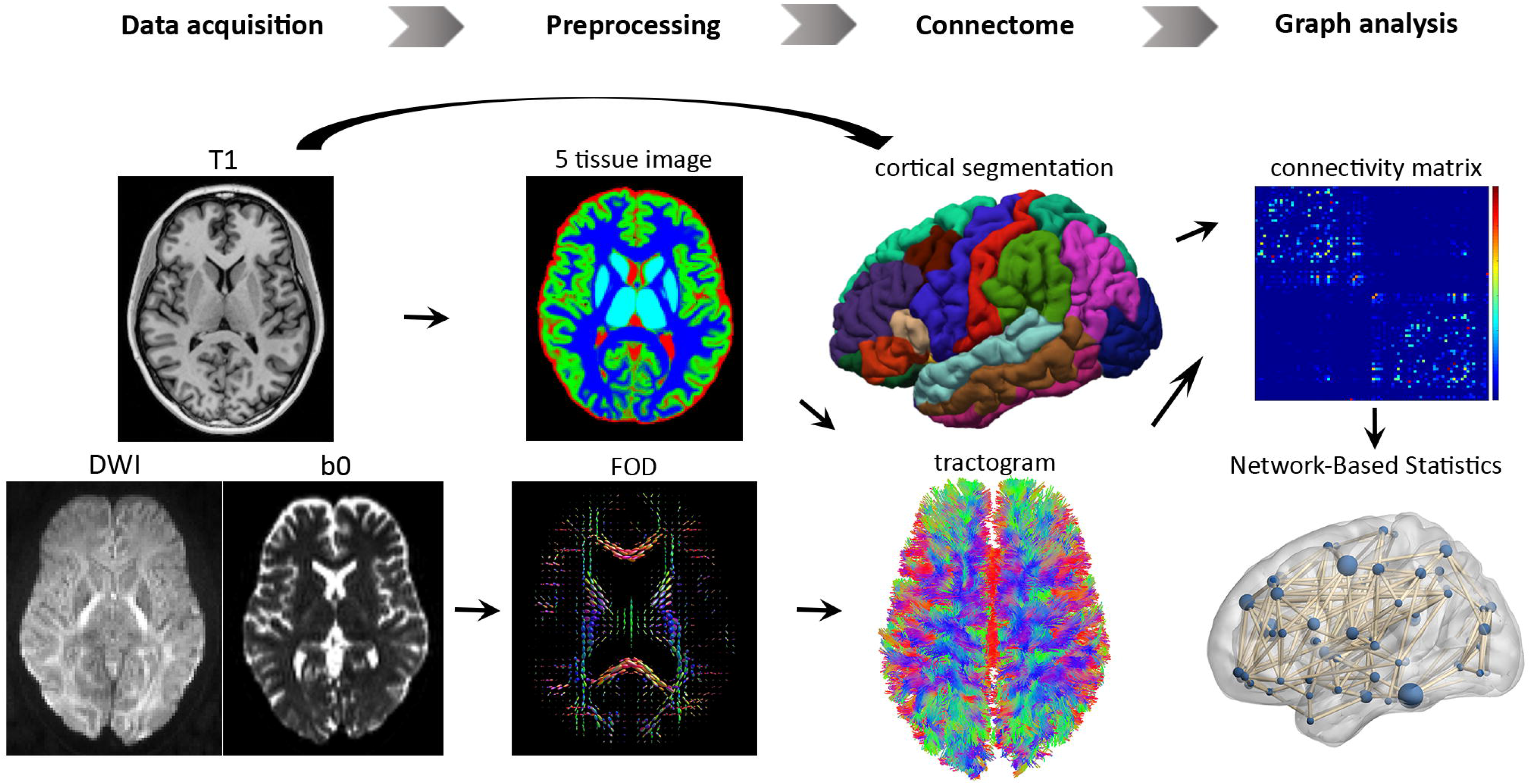
Flowchart of the steps involved in the structural connectome analysis. The T1-weighted image and diffusion weighted image (DWI) were co-registered, from which a five-tissue-type segmented image was generated. Multi-shell, multi-tissue response functions were estimated and multi-tissue constrained spherical deconvolution was applied to obtain fiber orientation distributions (FOD). Cortical segmentation was used to define 84 structural nodes and individual tractograms were generated for each participant using anatomically constrained tractography. Separate connectivity matrices were populated with the streamline count between the corresponding pairs of nodes, as well as fractional anisotropy (FA) estimates averaged over all voxels traversed by streamlines between node pairs. Network-based statistics were used to test for local white matter connectivity effects.

### 2.4 Statistical analysis

Network-based statistics (NBS) was used to investigate whether there was a linear relationship between PQ scores and white matter connectivity (Zalesky, Fornito, & Bullmore, 2010). For this purpose, two multiple linear regressions were employed with PQ scores as the predictor variable and the interregional connectivity matrices for streamline count and FA as outcome variables. Distress was added to the regression models as a nuisance covariate. Test statistics were computed independently for each connection. Supra-threshold connections were considered if their test statistic exceeded a *p*-value of .001 with 5000 permutations (t-statistic > 3). Subnetworks (connected components) were then identified with a family-wise error (FWE)-corrected *p*-value of .05.

## 3. Results

Participants’ scores on the PQ ranged from 0-162 (*M* = 49.34, *SD* = 43.99) and on distress from 0-48 (*M* = 11.59, *SD* = 13.35).

### 3.1 Connectivity disruptions

NBS identified a subnetwork of reducing streamline count with increasing PQ score (*p* = .048, FWE) (see Figure 2A). The network of significant connectivity reductions comprised 54 brain regions and 92 edges, affecting all cortical lobes, subcortical structures and the cerebellum. These results indicate that inter-regional connectivity strength declines with increasing number of psychotic-like experiences. A connectogram and a connectivity matrix were constructed to visualize the disrupted white matter network (see Figure 2B and 2C). Most connectivity abnormalities were identified between inter-hemispheric regions (58 edges) and in the right (25 edges) more so than the left (9 edges) hemisphere (see Figure 2B and 2C). Figure 2D shows a significant linear decline of mean streamline count in the identified subnetwork with increasing PQ scores (*r* = -.201, *p* = .0295), indicating that psychotic-like experiences increase with decreasing white matter connectivity. No significant subnetworks of reduced streamline count were observed for decreasing PQ scores, nor did we find reduced FA with increasing PQ.

**Figure 2.**
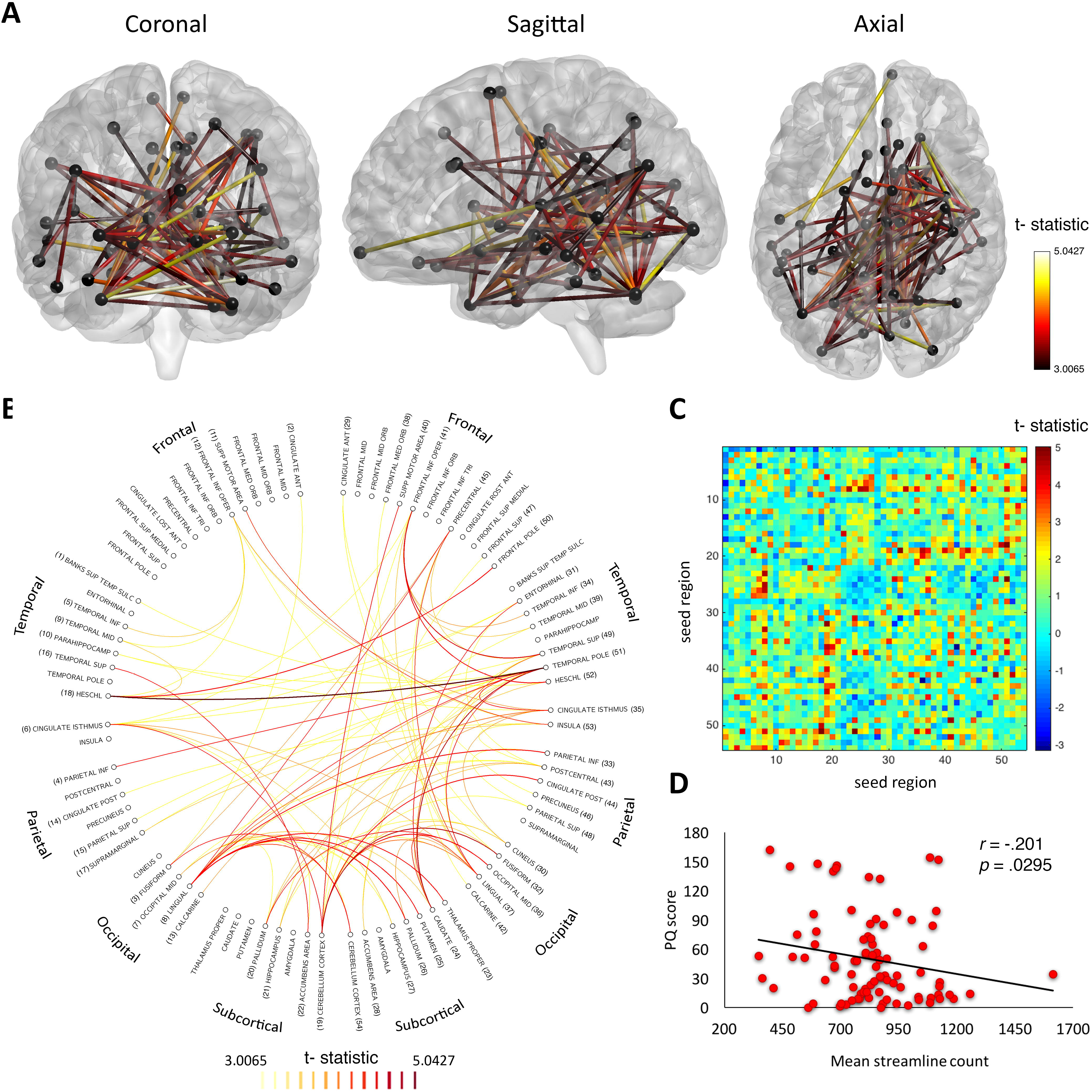
T-statistic is set to a threshold of 3, which corresponds to *p* = .001. Subnetworks are significant at a supra-threshold of *p_FWE_* = < .05. A) Widespread network of linearly declining streamline count with increasing number of prodromal psychotic experiences in healthy individuals. The network comprised 54 nodes and 92 edges. B) Connectogram of linearly declining streamline count with increasing PQ scores. Left hemisphere structures are represented on the left and right hemisphere structures on the right. Connection color represents t-statistic magnitude, ranging from 3 (yellow) to 5 (red). C) Connectivity matrix of the significant subnetwork consisting of 54 connected brain regions (seed region). D) Mean streamline count declines linearly with increasing PQ scores, indicating white matter connectivity loss with increasing psychotic-like experiences.

## 4. Discussion

Studies of white matter pathology in schizophrenia are plentiful, however the extent to which these changes translate to sub-clinical manifestations of psychotic-like experiences and whether white matter pathology in prodromal stages of psychosis could be used as biomarkers to identify individuals at high-risk for developing florid psychosis, remains unanswered. The aim of the present study was to investigate whether healthy individuals with a high quantity of psychotic-like experiences exhibit white matter disruptions in brain networks previously identified in patients with schizophrenia. As predicted, connectivity strength linearly declined with increasing number of psychotic-like experiences, similar to previous reports of connectivity loss from early to chronic stages of schizophrenia (Di Biase et al., 2017).

Similar to the present study, a study by Drakesmith et al. (2015) also investigated the structural connectome in healthy individuals with a high quantity of psychotic-like experiences but did not find any significant white matter network disruptions. These inconsistencies might be explained by the fact that the study by Drakesmith et al. (2015) used the psychotic-like symptoms semi-structured interview, which only assesses the positive symptoms hallucinations, delusions and bizarre symptoms (Horwood et al., 2008). Our study on the other hand, used the prodromal questionnaire (Loewy et al., 2005) which measures positive, negative and depressive symptoms. It has previously been reported that the prodromal phase of schizophrenia is marked by a variety of symptoms such as depression, fatigue, anxiety, psychotic symptoms, social withdrawal and irritability (Yung & McGorry, 1996; Yung et al., 2007). While the study by Drakesmith et al. (2015) investigates a group of heathy individuals with a high quantity of positive symptoms, the present study focuses on a more comprehensive set of prodromal psychotic-like experiences. It is therefore possible that the disrupted white matter network in the present study is associated with prodromal psychotic-like experiences in general rather than positive symptoms specifically.

Structural connectome studies have provided evidence that white matter pathology in schizophrenia is more widespread (Di Biase et al., 2017; Fornito et al., 2012; van den Heuvel et al., 2010; Wei et al., 2017; Zalesky et al., 2011) than previously anticipated from localized, regions of interest (ROI) approaches (Kubicki et al., 2007). The common finding of reduced fronto-temporal white matter connectivity in earlier studies of schizophrenia (Orrù, Pettersson-Yeo, Marquand, Sartori, & Mechelli, 2012) might be due to the fact the ROI approaches focus on specific, hypothesis-driven white matter pathways. Instead, our study adopted a whole brain approach which revealed a network of disrupted white matter connectivity across a number cortical and subcortical regions as opposed to white matter changes localized to fronto-temporal white matter pathways, which is in line with recent reports of widespread white matter reductions in schizophrenia (Pettersson-Yeo et al., 2011).

The findings of widespread white matter reductions in schizophrenia and in healthy individuals with psychotic-like experiences are in line with whole-brain changes reported from post-mortem, neuroimaging and histology studies in schizophrenia. Neuropathology studies report global reductions of neural density and a decrease of axons, dendrites and glia cells to be associated with schizophrenia (Boksa, 2012). These findings are supported by molecular studies, which report reductions in several presynaptic protein markers across the brain in patients with schizophrenia (Faludi & Mirnics, 2011). Post-mortem studies have provided evidence that several neurotransmitter systems across the brain are abnormal in schizophrenia, including dopamine, serotonin, γ-aminobutyric acid (GABA) and glutamate (Harrison, 2000). Furthermore, neuroimaging studies have reported widespread reduced cortical folding in patients with schizophrenia, which has been interpreted to reflect abnormal neural connectivity during brain maturation (Sallet et al., 2003). These findings, together with the findings from the present study, indicate that brain changes related to schizophrenia are more widespread than initially anticipated and that whole-brain approaches might be more suitable to study the pathophysiology of schizophrenia as opposed to ROI approaches.

An ongoing limitation of structural connectomics studies is the large discrepancies in acquisition parameters and data analysis. Contrary to many previous structural connectomics studies, which used diffusion tensor imaging, the present study used a multi-shell, high angular resolution diffusion imaging acquisition protocol, which can provide information about crossing fibers on the sub-voxel level. This makes the comparison of connectomics studies difficult and calls for a standardization of acquisition protocols and analyses pipelines. Another limitation of the present study is the lack of a clinical schizophrenia group. A direct comparison of structural connectomics between healthy individuals with a high quantity of psychotic-like experiences and schizophrenia patients would provide a better understanding of brain networks that might be particularly useful to investigate as potential prodromal psychosis biomarkers and therefore, we suggest, provides a fruitful avenue for future research.

In sum, we identified a widespread network of white matter connections that declined in streamline count as psychotic-like experiences in healthy individuals increased, similar to the white matter connectivity reductions reported from early to chronic stages of schizophrenia (Di Biase et al., 2017). The disrupted network spanned over several cortical and subcortical brain regions, which is in line with findings from post-mortem, neuroimaging and molecular studies of schizophrenia that report abnormalities in schizophrenia across the entire brain as opposed to circumscribed regions. The findings from this study indicate that widespread white matter disruptions are already apparent in nonclinical individuals who display psychotic-like experiences. We speculate that such white matter alterations may be a potential biomarker to identify individuals at high-risk for transitioning to florid psychosis, who might benefit from preventative treatments.

## Acknowledgments

This work was funded by the Australian Research Council Centre of Excellence for Integrative Brain Function (ARC Centre Grant CE140100007), a University of Queensland Fellowship (2016000071), and a Foundation Research Excellence Award (2016001844) to MIG, as well as a University of Queensland International Research Scholarship to RR. The authors thank the participants for their time and Aiman Al-Najjar and Nicole Atcheson for assisting with data collection.

## Conflict of interest

The authors declare no competing financial interests

